# A comprehensive knowledge base of synaptic electrophysiology in the rodent hippocampal formation

**DOI:** 10.1101/632760

**Authors:** Keivan Moradi, Giorgio A. Ascoli

**Affiliations:** Neuroscience Program, Krasnow Institute for Advanced Study, George Mason University, Fairfax, VA (USA); Bioengineering Department, Krasnow Institute for Advanced Study, George Mason University, Fairfax, VA (USA)

**Keywords:** Synapses/physiology, Neuron types, Knowledge Bases, Information Storage and Retrieval, Circuit biophysics, Computational Biology, Models

## Abstract

The cellular and synaptic architecture of the rodent hippocampus has been described in thousands of peer-reviewed publications. However, no human- or machine-readable public catalog of synaptic electrophysiology data exists for this or any other neural system. Harnessing state of the art information technology, we have developed a cloud-based toolset for identifying empirical evidence from the scientific literature pertaining to synaptic electrophysiology, for extracting the experimental data of interest, and for linking each entry to relevant text or figure excerpts. Mining more than 1,200 published journal articles, we have identified eight different signal modalities quantified by 68 different methods to measure synaptic amplitude, kinetics, and plasticity in hippocampal neurons. We have designed a data structure that both reflects these variabilities and maintains the existing relations among experimental modalities. Moreover, we mapped every annotated experiment to identified “synapse types,” i.e. specific pairs of presynaptic and postsynaptic neuron types. To this aim, we leveraged Hippocampome.org, an open-access knowledge base of morphologically, electrophysiologically, and molecularly characterized neuron types in the rodent hippocampal formation. Specifically, we have implemented a computational pipeline to systematically translate neuron type properties into formal queries in order to find all compatible synapse types. With this system, we have collected nearly 40,000 synaptic data entities covering 88% of the 3,120 potential connections in Hippocampome.org. Correcting membrane potentials with respect to liquid junction potentials significantly reduced the difference between theoretical and experimental reversal potentials, thereby enabling the accurate conversion of all synaptic amplitudes to conductance. This dataset allows for large-scale hypothesis testing of the general rules governing synaptic signals. To illustrate these applications, we confirmed several expected correlations between synaptic measurements and their covariates while suggesting previously unreported ones. We release all data open source at Hippocampome.org in order to further research across disciplines.

## Introduction

Synaptic communication is essential for understanding the genesis, dynamics, and function of neuronal ensembles. The electrophysiology of synapses is often characterized in terms of signal amplitude, kinetics (delay and duration or rise and decay time), and short- or long-term plasticity. Each of these characteristics depends on the combined properties of the pre- and post-synaptic neurons and varies widely across and within neural systems. The rodent hippocampus has long served as a discovery sandbox for synaptic biophysics (Buzsaki, 1984; Freund & Buzsaki, 1996; Nicoll, 2017; Pelkey et al., 2017). Both excitatory and inhibitory synapses in the hippocampal formation exhibit tremendous diversity in a number of mechanisms including synchronous or asynchronous release (Daw, Tricoire, Erdelyi, Szabo, & McBain, 2009; Szabo, Holderith, Gulyas, Freund, & Hajos, 2010; Szabo, Papp, Mate, Szabo, & Hajos, 2014), failure rate (Losonczy, Biro, & Nusser, 2004; Maccaferri, Roberts, Szucs, Cottingham, & Somogyi, 2000), and potentiation or depression (Alle, Jonas, & Geiger, 2001; Jappy, Valiullina, Draguhn, & Rozov, 2016).

Despite the rich publication history in this field, no systematic data mining study has so far catalogued the properties of these synapses. Thus, a consistent synaptic inventory of different electrophysiological variants has yet to be made available online for the rodent hippocampus or any other neural circuit. Quantitative information about synapses is valuable for experimental scientists to ensure proper replicability of results, to identify knowledge gaps, and to enable congruent comparison of data. Detailed knowledge of synaptic properties allows computational neuroscientists to constrain and validate increasingly predictive simulations (Markram et al., 2015). A systematic and consistent knowledge base may also produce new discovery opportunities for data scientists in the spirit of ongoing large-scale, collaborative, and multinational efforts such as the BRAIN initiative and the Human Brain Project (Kandel, Markram, Matthews, Yuste, & Koch, 2013). Moreover, in light of recent progress in biologically inspired artificial intelligence (Ullman, 2019) and the design of neuromorphic chips empowered with artificial synapses (Wan, 2018; L. Q. Zhu, Wan, Guo, Shi, & Wan, 2014), both the scientific community and different industries have a growing interest in synaptic data to fuel data-driven modeling endeavors and the incubation of new technologies.

The Hippocampome.org project effectively organized a vast amount of information on the rodent hippocampal microcircuit at the cell type level (Wheeler et al., 2015). This knowledge base identifies 122 neuron types across the dentate gyrus, areas CA3, CA2, CA1, subiculum, and entorhinal cortex based on their main neurotransmitter released (glutamate or GABA), the laminar distribution of axons and dendrites, and converging molecular and electrophysiological evidence. This framework conveniently extends to the notion of “synapse type,” defined as a directional potential connection in a unique pair of presynaptic and postsynaptic neuron types (Rees et al., 2016). Based on available evidence regarding neuronal morphology and known targeting specificities (Rees, Moradi, & Ascoli, 2017), we estimate the existence of at least 3,120 potential connections in the hippocampal formation (Hippocampome.org/connectivity). For how many of these synapse types are any experimental measurements available concerning at a minimum signal strength, time course, and plasticity?

Conducting a methodical data mining study for synaptic electrophysiology constitutes a formidable challenge. On the one hand, inconsistent terminology of neuron names and properties (Hamilton et al., 2017) renders fully automated text-mining approaches unreliable. On the other, purely manual data extraction and annotation from the scientific literature are excruciatingly labor intensive, time consuming, and error prone, because of the difficulties of unambiguously determining cell type based on morphological features and of detecting synaptic signal in published figures. In particular, most electrophysiological studies adopt their own custom definition of neuron types, often based on pragmatic requirements or limitations of the experimental design. Therefore, neuron groups from each research report can be typically mapped onto multiple distinct Hippocampome.org types. Furthermore, the definitions of data modalities and of measured values are also often inconsistent in peer-reviewed publications.

In order to tackle the above challenges we have recently devised a systematic workflow combining state-of-the-art information technology with carefully vetted domain expertise (Moradi & Ascoli, 2019). We split each relevant scientific article into several unique experiments delimiting finite sets of synapse types. We then translate the morphology, molecular markers, membrane electrophysiology, and firing patterns of the possible pre- and post-synaptic neurons into machine-readable queries. We have built a search engine to translate each query into a dynamic list of potential connections. In parallel, the synaptic electrophysiology data, either in the body or the figures of each paper, are semi-automatically annotated, extracted, digitized, and linked to proper references and comprehensive metadata using a custom-designed cloud-based data mining toolset.

Here we present quantitative synaptic electrophysiology results using the above described data processing pipeline. Specifically, with this work we publicly release over 8,000 pieces of annotated experimental evidence from more than 1,200 journal articles, accounting for nearly 90% of synapse types in the rodent hippocampal formation. We demonstrate the richness of these data by reconciling the observed and theoretical reversal potential values for the main excitatory and inhibitory ionotropic receptors (AMPA and GABA_A_) after correction for liquid junction potential, by quantifying known interactions between recorded synaptic parameters and common experimental covariates, and by reporting several novel correlations for unitary GABAergic currents.

## Methods

### Inclusion criteria and literature search methodology

The scope of this work concerns the monosynaptic electrophysiology of non-cultured, healthy, adult or young adult (>P12) rodent hippocampal formation: dentate gyrus, CA3, CA2, CA1, subiculum, and entorhinal cortex. The data sources included all peer-reviewed original articles containing direct experimental evidence pertaining to synaptic signals (i.e., excluding reviews and book chapters). The relevant corpus was collated in three steps. We started by mining the 476 papers already included in the bibliography of Hippocampome.org v1.3 that met these criteria. Next, we searched all references of and citations to those papers through 2018. We assessed each new article for the presence of measurements of synaptic signals accompanied by relevant information to identify the corresponding neuron types, including morphology, molecular expression, and intrinsic electrophysiology. We annotated every article with pertinent content, lack thereof or the reason for exclusion. Lastly, we performed literature searches specifically targeted at all potential connections still devoid of synaptic information. We also set a PubMed alert for the query ‘interneuron AND (hippocampus OR CA1 OR CA2 OR CA3 OR “dentate gyrus” OR subiculum OR “entorhinal cortex”) AND (IPSP OR IPSC OR EPSP OR EPSC)’ to maintain the knowledge base updated with forthcoming publications.

### Data mining procedures

We have implemented a set of cloud-based tools to assist with the critical aspects of data mining. We used Google Apps Script cloud computing framework to develop the backend and CSS, HTML5, and JavaScript to design the frontend. To encourage collaboration and data re-usage, we freely release with this study all mined data, tools, source codes, and users’ manuals via Hippocampome.org/synaptome.

#### Text analysis

The central elements of interest in each relevant article are the reported synaptic signals in the form of either recorded traces or quantitative measurements of amplitude, kinetics, and plasticity. Besides synaptic signals, we also identified in every article any figure or text excerpt corresponding to three distinct types of additional content: properties that may define neuron types, experimental metadata, and other useful information. Pertinent neuron type properties include anatomy (e.g. somatic location and laminar distribution of axons), molecular biomarkers (e.g. expression of parvalbumin and lack of somatostatin), membrane electrophysiology (e.g. input resistance and time constant), and firing patterns (e.g. rapidly adapting or persistent bursting). Each of these characteristics may refer to, and are separately annotated for, presynaptic or postsynaptic neurons. Metadata consists of any covariates that may change the properties of synaptic signals, such as animal age and strain, drugs, solutions, temperature, and recording conditions and settings. Other useful information includes, among many others, numerical ratios of different neuron types, evidence of synaptic specificity, and connection probabilities.

#### Informatics tools

In order to make the systematic extraction of the above details less error-prone, time-consuming, and labor-intensive, we have custom designed a dedicated Google Sheets graphical user interface (“Review” function at Hippocampome.org/synaptome). This tool is ergonomically optimized to assist in metadata annotation and automatically highlights potential areas of interest in the text excerpts. An accompanying “Text Cleaner” tool prepares the excerpts for editing and labeling by autonomously standardizing and correcting all frequent formatting inconsistencies (special characters, Greek letters, symbols, etc.). The “data extraction” tool dynamically presents to the user a series of fillable forms with fields relevant to the identified measurement methods after pre-compiling any suitable entry with information already recognized in the text annotation phase. Lastly, a “Check Query” tool maps each experiment to its appropriate subset of Hippocampome.org potential connections (Hippocampome.org/connectivity). This is achieved by translating the neuron type properties of each experiment into a custom machine-readable query (Hippocampome.org/query) and by calling via web API a PHP search engine to match those properties to the corresponding potential connections.

### Data quantification

In order to assess quantitatively the amount of the knowledge base diverse content, we adopt and extend the terminology utilized by Hippocampome.org (Wheeler et al., 2015). For the purpose of this study, a **piece of evidence** (PoE) is a numerical or categorical data entity that describes a synaptic signal. While categorical data are typically extracted directly from papers, numerical data can also be obtained by quantifying digitized synaptic traces. Any independent measurement or observation generally constitutes a distinct PoE. For example, two recordings of the same neuron pair at different extracellular calcium concentrations will produce two PoEs. All numerical or categorical data entities that describe experimental conditions and covariates rather than synaptic signals are considered **metadata** (Table 1). Numerical data are typically expressed as combinations of central tendency (mean or median) and variance (standard error, standard deviation or interquartile range) or else a range (lower and upper limits), plus a sample size. In other words, a single numerical PoE or covariate may consist of up to five values to describe a sample distribution, plus eventual text comments.

**Table 1:**
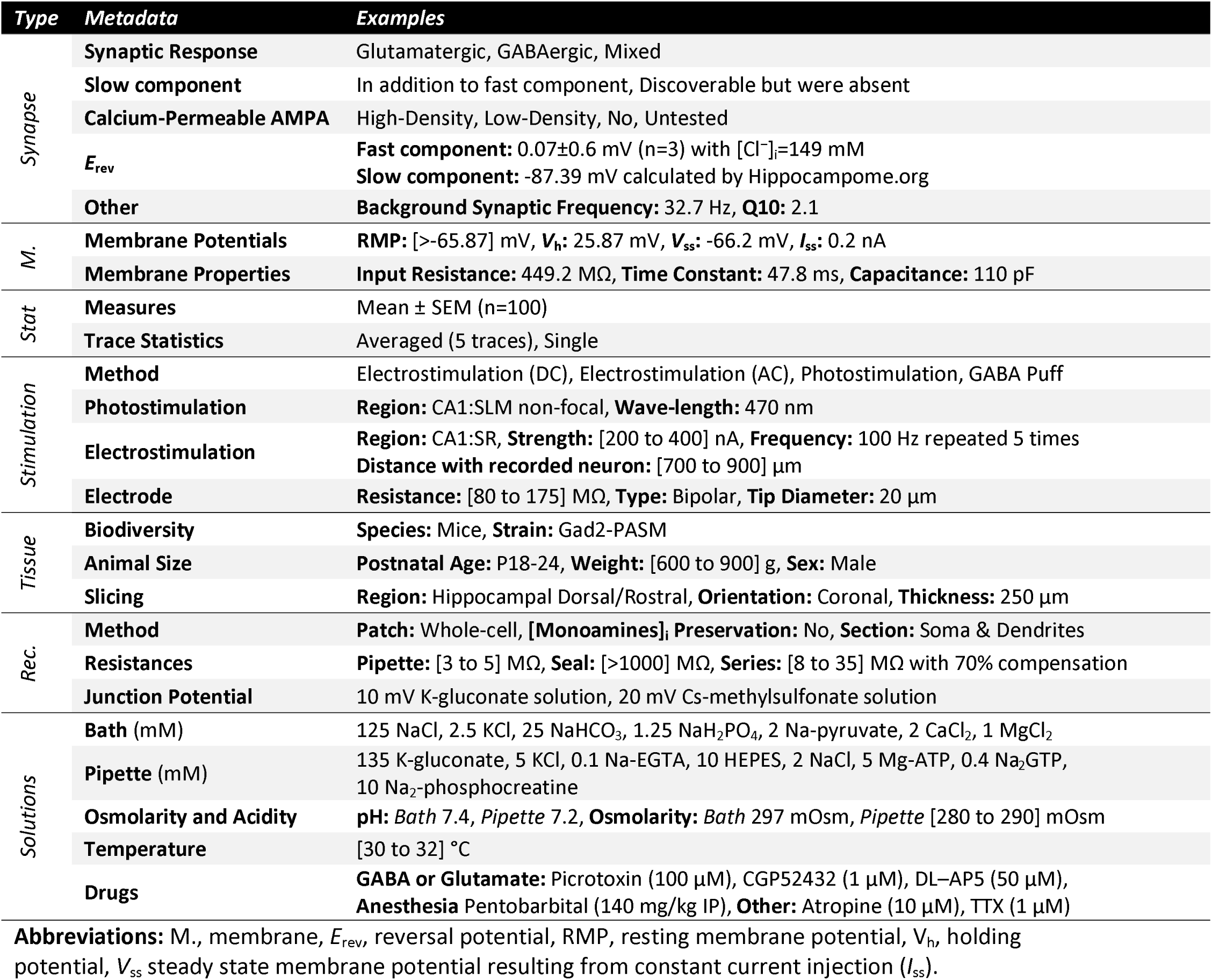
Metadata list.

A **piece of knowledge** (PoK) is a conceptualized value or range of values for a specific synaptic property of a given synapse type: in other words, a parameter describing amplitude, kinetics, or plasticity for a unique pair or presynaptic and postsynaptic neurons that is supported by at least one PoE. For a particular neuron pair, different signal modalities and/or different measurements of the same specific synaptic property (e.g. 10-90% and 20-80% rise times) contribute to one and the same PoK (in this case, rise time for that synapse type). However, if the same PoE can be mapped onto six synapse types (e.g. any combination of two presynaptic neuron types and three postsynaptic neuron types), it will produce six PoKs.

### Data normalization and analysis

We implemented a reversal potential (*E*_rev_) and recording pipette liquid junction potential calculator in JavaScript running on the Google Cloud.

#### Reversal potential calculation

The code first calculates the intracellular ionic concentrations from the pipette solution content based on recording method. For whole-cell and outside-out modalities, the intracellular concentrations are set to be equal to the pipette concentrations. For cell-attached experiments, in contrast, the following standard textbook (Hille, 2001) ionic concentrations are assumed (in mM): [Na^+^]_i_=10, [K^+^]_i_=140, [Cl^−^]_i_=4, [Ca^2+^]_i_=10^−4^, [Mg^2+^]_i_=0.8, and [HCO_3_^−^]_i_=12. For sharp-electrode recordings, we empirically assume 1% ion exchange relative to the concentration difference at the start of the experiment,

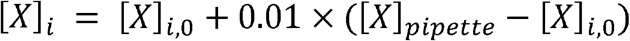

where *X* is the ion, *[X]*_*pipette*_ is the ionic concentration in the pipette, and *[X]*_*i,0*_ is the intracellular concentration before cell impalement. The same 1% ionic exchange also applied to the perforated-patch method, but only for permeable monovalent cations (Finkelstein & Andersen, 1981). Next, the algorithm corrects the concentrations by considering the effect of weak acids like HEPES and polyvalent chelating agents like EGTA that do not fully ionize in solution, and converts all resulting values to ionic activities (Davies, 1938). Finally, the calculator derives the reversal potential from ionic activities solving numerically the Goldman-Hodgkin-Katz current equation for channels permeable to ions with any number of valences (Hille, 2001). Since ionic valences for synaptic channels in the scope of this study are limited to +1, −1, or +2, we employed Lewis’ voltage equation (Lewis, 1979) for efficient initialization of the numerical solver (Loisel, 2012).

For the relative ionic permeability parameters in the Goldman-Hodgkin-Katz equation, we used available experimentally estimates for AMPA, NMDA, GABA_A_ and GABA_B_ channels (Farrant & Kaila, 2007; Jatzke, Watanabe, & Wollmuth, 2002; Traynelis et al., 2010). The GABA_A_ channel is considered permeable to Cl^−^, HCO_3_^−^, Br^−^, and F^−^, as well as to gluconate if explicitly noted. For GABA_B_, we included both K^+^ and Na^+^ with an empirically determined ratio P_Na_/P_K_= 0.02, compatible with earlier research (Hille, 1973; Sah, Gibb, & Gage, 1988). Calciumim-permeable AMPA channel were presumed to be permeable to Na^+^, K^+^, and Cs^+^. Calcium-permeable AMPA and NMDA channels included the same ionic species plus Ca^2+^. The relative calcium permeability of AMPA depends on membrane potential, increasing from 0.6 at −60 mV to 0.92 at −20 mV (Jatzke et al., 2002) and we assumed a linear function.

#### Junction potential calculation

The junction potential (*V*_j_) is observed at the tip of the recording electrode, where ion exchange occurs between the pipette solution and the bath or the intracellular solutions. Faster moving ions leave behind slower ions that may have opposite charge. These ionic mobility differences lead to an electric field at the interface, or junction, between the two liquids. The generalized Henderson equation derives *V*_j_ from ionic activities and experimentally determined ion mobilities (Barry, 1994; Barry & Lynch, 1991; Marino, Misuri, & Brogioli, 2014; Morf, 2012). We correct *V*_j_ between the extracellular and pipette solutions for all recording methods, except for the sharp-electrode, for which we additionally correct for the difference of *V*_*j*_ between the pipette and intracellular solutions. The calculator automatically chooses the appropriate *V*_*j*_ sign for current- or voltage-clamp recordings. If an article reports the experimental *V*_*j*_ measurement, we use the value, but choose the sign based on the calculated *V*_*j*_.

#### Statistics and illustrations

For correlation analysis and figure plotting we used publicly available R packages including R Markdown, Tidyverse, ggpubr, Venneuler, and UpSetR (Baumer, Cetinkaya-Rundel, Bray, Loi, & Horton, 2014; Lex, Gehlenborg, Strobelt, Vuillemot, & Pfister, 2014; Wickham, 2016, 2017; Wilkinson & Urbanek, 2011). We used Pearson and Spearman coefficients as measures of linear and nonlinear correlations, respectively. We assessed statistical differences with Wilcoxon non-parametric test and considered results significant when P < 0.05 after “False Discovery Rate” multiple-testing correction (Benjamini & Hochberg, 1995).

## Results

### Mapping Experiments to Synapse Types

Any unique combination of potentially connected presynaptic and postsynaptic neuron types produces a specific synapse type. Consider as an example a simple case of inhibitory convergence on the principal cells of the dentate gyrus (Fig. 1). This minimalist circuit consists of three different neuron types: granule as postsynaptic cell and two GABAergic interneurons as presynaptic cells, for instance hilar commissural pathway associated (HICAP) and basket neurons. The potential connection between HICAP (a dendritic-targeting, non-fast-spiking interneuron type) and granule cells is produced by the co-localization of the pre-synaptic axon and postsynaptic dendrites in the inner molecular layer (Fig. 1A). Similarly, the synapse type between (perisomatic-targeting, fast-spiking) basket and granule cells is consistent with the spatial overlap of the presynaptic axon with the postsynaptic soma and proximal dendrites in the granular layer. Even though individual synaptic traces might not represent the population averages, the signals recorded from these two pairs display evidently distinct amplitudes and kinetics (Fig. 1B). Conversely, granule cells also elicit diversified synaptic responses in different postsynaptic interneuron types (Toth, Suares, Lawrence, Philips-Tansey, & McBain, 2000).

**Figure 1.**
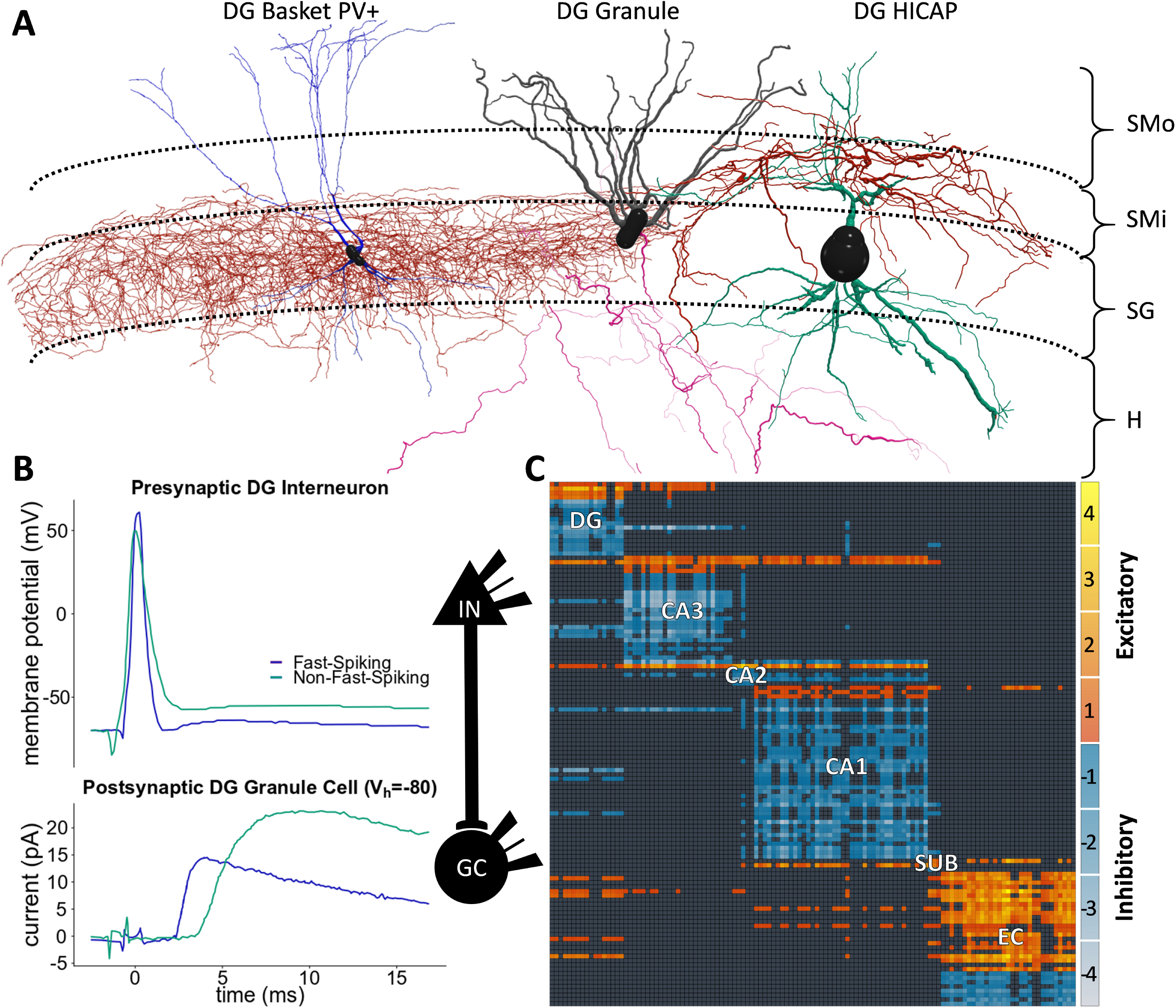
From neuron types to synapse types. **(A)** Red, the axons of the dentate gyrus (DG) basket neuron type innervate SG (stratum granulare), and those of the HICAP type invade SMi (stratum moleculare – inner). Blue and green, the dendrites of both neuron types span all four DG layers. The local axons of granule cells innervate the hilus (H) while its dendrites span SMi and SMo (stratum moleculare – outer). HICAP axons and granule dendrites are co-located, as are the basket axons and the granule perisomatic region; therefore, these neurons give rise to two distinct inhibitory synapse types. Morphologies rendered with neuTube (Feng, Zhao, & Kim, 2015) using data from the Bausch and Lien archives (Bausch, He, Petrova, Wang, & McNamara, 2006; Liu, Cheng, & Lien, 2014) of NeuroMorpho.Org (Ascoli, Donohue, & Halavi, 2007). **(B)** The presynaptic spikes (upper traces) generate postsynaptic signals (lower traces) digitized and plotted from pair recording (Liu et al., 2014), from which we identified “Fast-Spiking” as DG basket and “Non-Fast-Spiking” as DG HICAP (IN: interneuron). **(C)** The 122 known neuron types in the rodent hippocampal formation (presynaptic: rows; postsynaptic: columns) form 3,120 synapse types. The heat map (SUB: subiculum; EC: entorhinal cortex) represents the number of distinct layers in which excitatory and inhibitory axons co-localize with relevant postsynaptic elements. For instance, the inhibitory synapse types in (A) have only one co-location each (in SMi and SG, respectively), corresponding to a −1 value in the directional connectivity matrix.

The magnitude of the problem begins to become apparent when considering that the 122 known neuron types in the rodent hippocampal formation generate as many as 3,120 distinct synapse types (Fig. 1C). This intrinsic circuit complexity is cumulated with another challenge: the uncertain identification of synapse types in available experimental reports. The above example clearly pinpoints individual presynaptic as well as postsynaptic neuron types and therefore unique synapse types, but this is hardly typical. More often studies do not report the reconstructed morphologies of the stimulated or recorded neurons, but instead simply describe the common properties of different neuron types. If multiple neuron types share these properties, mapping the reported data to a single synapse type becomes impossible. Even the most accurate paired-recording experiments, in which all cell types are adequately reconstructed, may suffer from result pooling. Nonetheless, it is usually possible to map a given experimental description to a limited set of synapse types.

Different data entities from the same publication may map to distinct sets of synapse types. The same study illustrated above, for example, also grouped many stimulated neuron types with diverse morphologies based on their spiking frequencies and soma location. Thus, our data mining process parses each paper into separate sections or “experiment” based on the common sets of identifiable neuron types (Fig. 2). For instance, a change in the stimulation region of the slice produces a distinct mapping, but a change in the extracellular calcium concentration does not. As described in the Methods, the properties of presynaptic and postsynaptic neurons are then mapped onto a specific group of synapse types. We consider a mapping as “proper” only if the corresponding experiment matches a single synapse type. For all other “fuzzy” mappings that involve multiple synapse types, we assign high and low confidence to the more and less likely neuron type pair(s), respectively. This determination relies on cell-type ratios, connection selectivity, or explicit assumptions of the original authors. In the experiment reported in Fig. 2, for instance, the HICAP-granule mapping is high-confidence due to the largest reported HICAP ratio among possible presynaptic types. Note that, although this work defines synapse types based on potential rather than established contacts, proper mappings are especially valuable because they also demonstrate the existence of (i.e. they “validate”) the corresponding connection.

**Figure 2.**
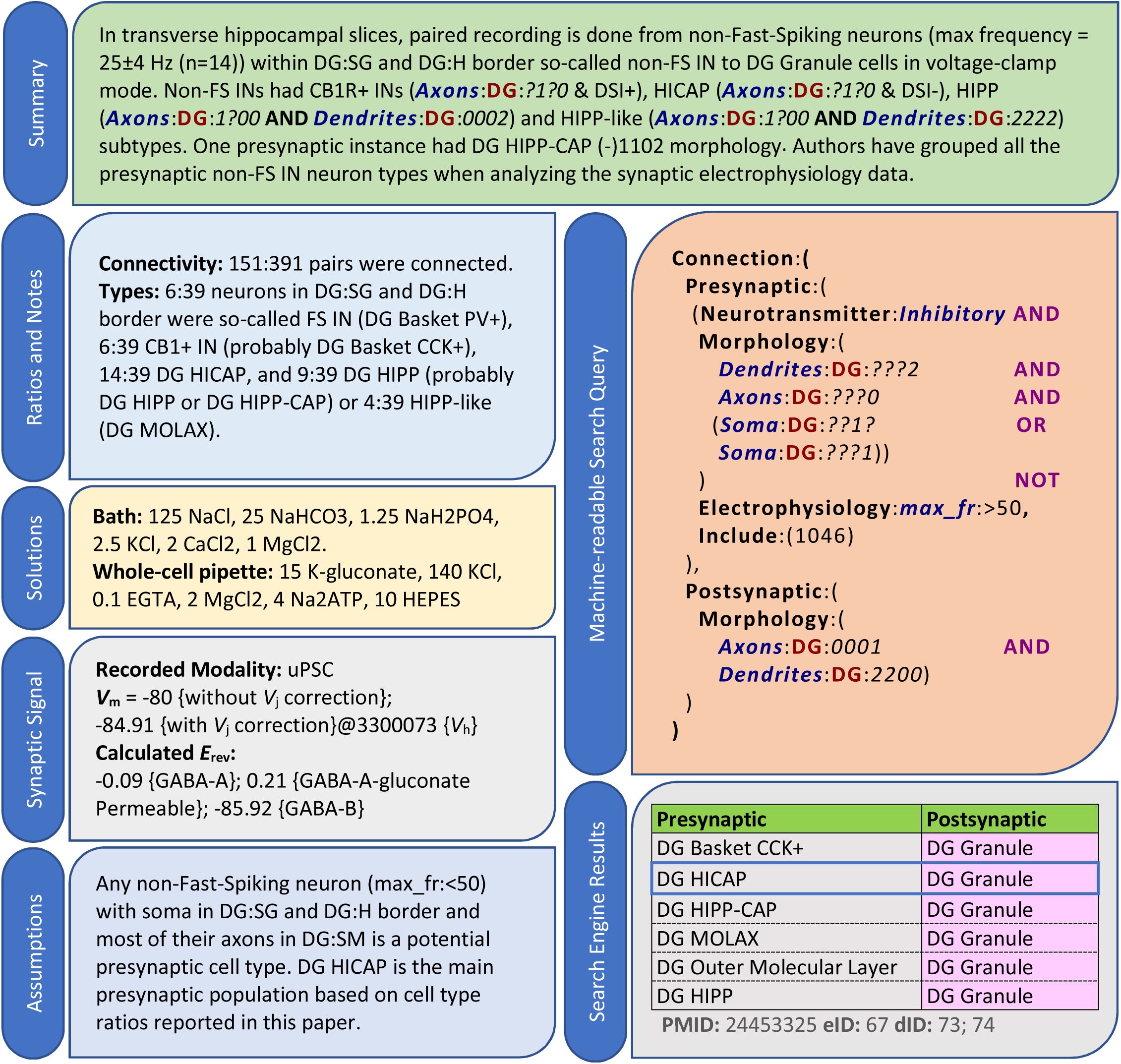
Literature mining and knowledge extraction. For every experiment, we provide (i) a summary; (ii) connectivity ratios, cell-types counts, and any other relevant notes; (iii) bath and pipette solutions; (iv) recorded modalities and pertinent data such as postsynaptic potential (*V*_m_), liquid junction correction (*V*_j_), and measured or calculated reversal potentials (*E*_rev_), each tagged with a reference ID; (v) needed assumptions for neuron identification; (vi) a machine-readable query; and (vii) mapped synapse types and related confidences (blue border: high confidence; others: low confidence), along with identifiers for the publication (PMID), experiment (eID), and extracted data IDs (dIDs). The data in this example is from (Liu et al., 2014).

### Dense Coverage of Synaptic Knowledge

Starting from 1,203 publications, we have extracted, annotated, and mapped the properties of 2,746 synapse types, or 88% of the 3,120 potential connections in the rodent hippocampal formation (Fig. 3). This proportion is remarkably stable across all major sub-regions of the hippocampal formation: 85.3% in dentate gyrus, 84.6% in CA3, 92.7% in CA1, 87.0% in entorhinal cortex, and 86.9% for projection synapses between sub-regions. Despite the richness of the hippocampus literature, only a minority (10.9%) of synapse types had at least one proper (n=71) or high-confidence fuzzy (n=229) mapping. Again, this proportion was essentially constant across CA3 (11.5%), CA1 (10.8%), and sub-region projections (10.9%), but was higher in dentate gyrus (17.7%) and lower in entorhinal cortex (5.6%). Out of over 3,000 potential connections, available experimental evidence can firmly validate merely 194 synapse types (Fig. 3A, grid pattern), including 123 based on electron microscopy (69 high-confidence fuzzy, 52 low-confidence fuzzy, 2 with no synaptic electrophysiology). Most synapse types with proper mapping also had high-confidence fuzzy mappings, and low-confidence fuzzy evidence was available for all of the high-confidence or properly-mapped types (Fig. 3B). The available data allow the determination of the amplitude and kinetics of most synapse types; plasticity information is available for slightly more than half of the cases, while other measurements, such as transmission failure and quantal release, for less than one third (Fig. 3C).

**Figure 3.**
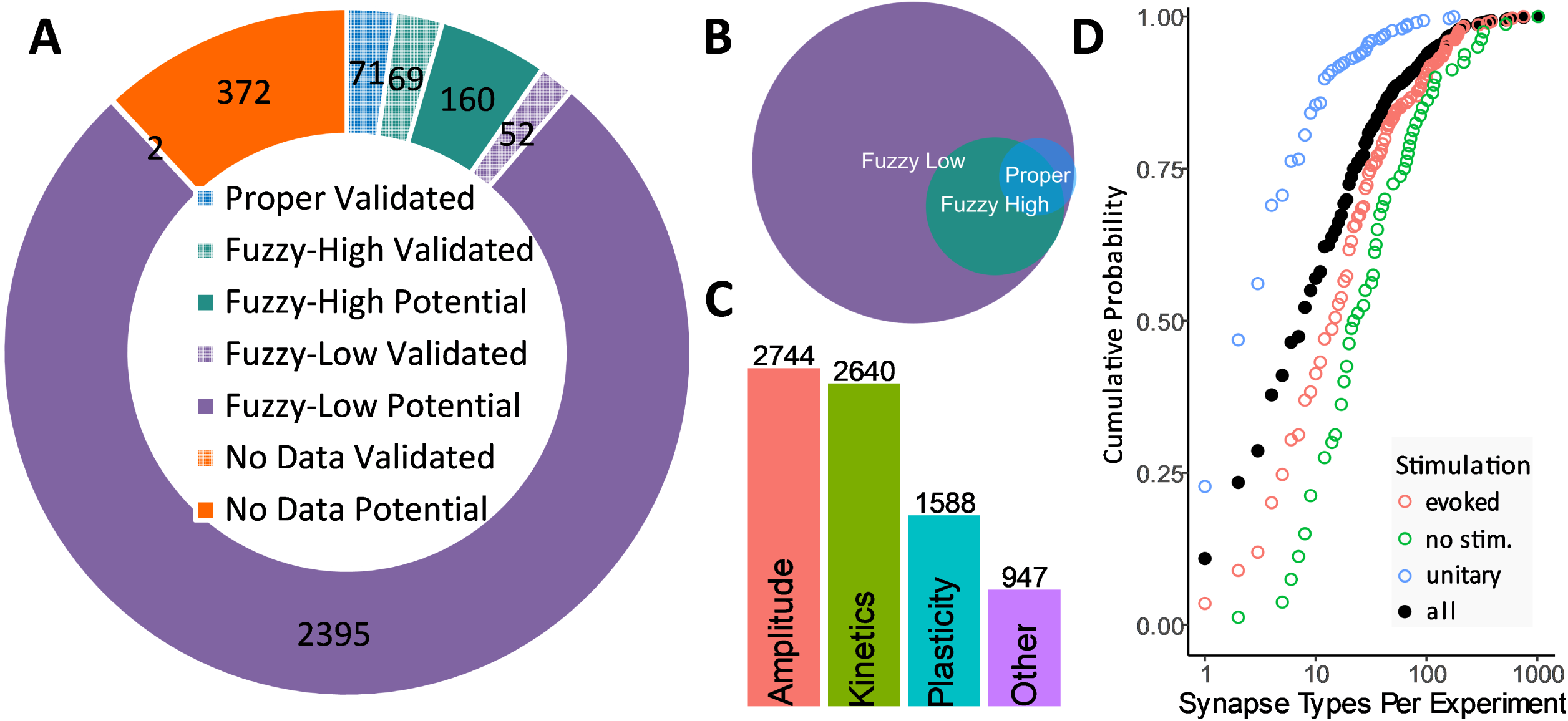
Mapping summary. **(A)** Integrated knowledge mapping (clockwise from top): “proper” (blue), “high-confidence fuzzy” (green), and “low-confidence fuzzy” (purple). Grid patterns indicate validated (as opposed to potential) connections. **(B)** An individual synapse type may be linked to multiple experiments with variable mapping confidence. **(C)** Amplitude and kinetics are the most prevalently reported synaptic electrophysiology properties. **(D)** Mapping degeneracy by stimulation method: unitary signals (mostly paired recording), evoked (extracellular) and spontaneous. Filled circles represent all methods together.

The extracted signals encompass four mechanism of synaptic activation: (i) “unitary,” resulting from the stimulation of an individual presynaptic neuron, as in paired-recordings; (ii) “evoked,” resulting from the stimulation of a population of presynaptic neurons, as in extracellular electro-, photo- or chemo-stimulation; (iii) “spontaneous,” reflecting background synaptic activity in the absence of stimulations controlled by the experimenter; and (iv) “miniature,” corresponding to unprompted neurotransmitter release while blocking action potentials. The choice of stimulation method greatly impacts mapping resolution: half of paired recordings but only 24% of all experiments are mapped to just one or two synapse types; due to the higher mapping degeneracy of evoked stimulation and (especially) spontaneous or miniature activity, the overall median number of synapse types per experiment is eight (Fig. 3D).

### Recording Modalities, Measurement Methods, and User Access

Postsynaptic signals are recorded as either potentials in current-clamp or currents in voltage-clamp, yielding eight different combinations with the four above-described stimulation methods (Fig. 4A). The most common modality is evoked current, but different modalities are often related in the same experiments, for instance when an investigator tests different stimulation methods and recording modes while clamping a postsynaptic neuron. Preserving the link between these data is particularly useful to integrate disparate data sources into a consistent analysis framework. The most common relation is between evoked currents and potentials (Fig. 4B), but a substantial number (>30) of experiments also offers valuable relations among three data modalities.

**Figure 4.**
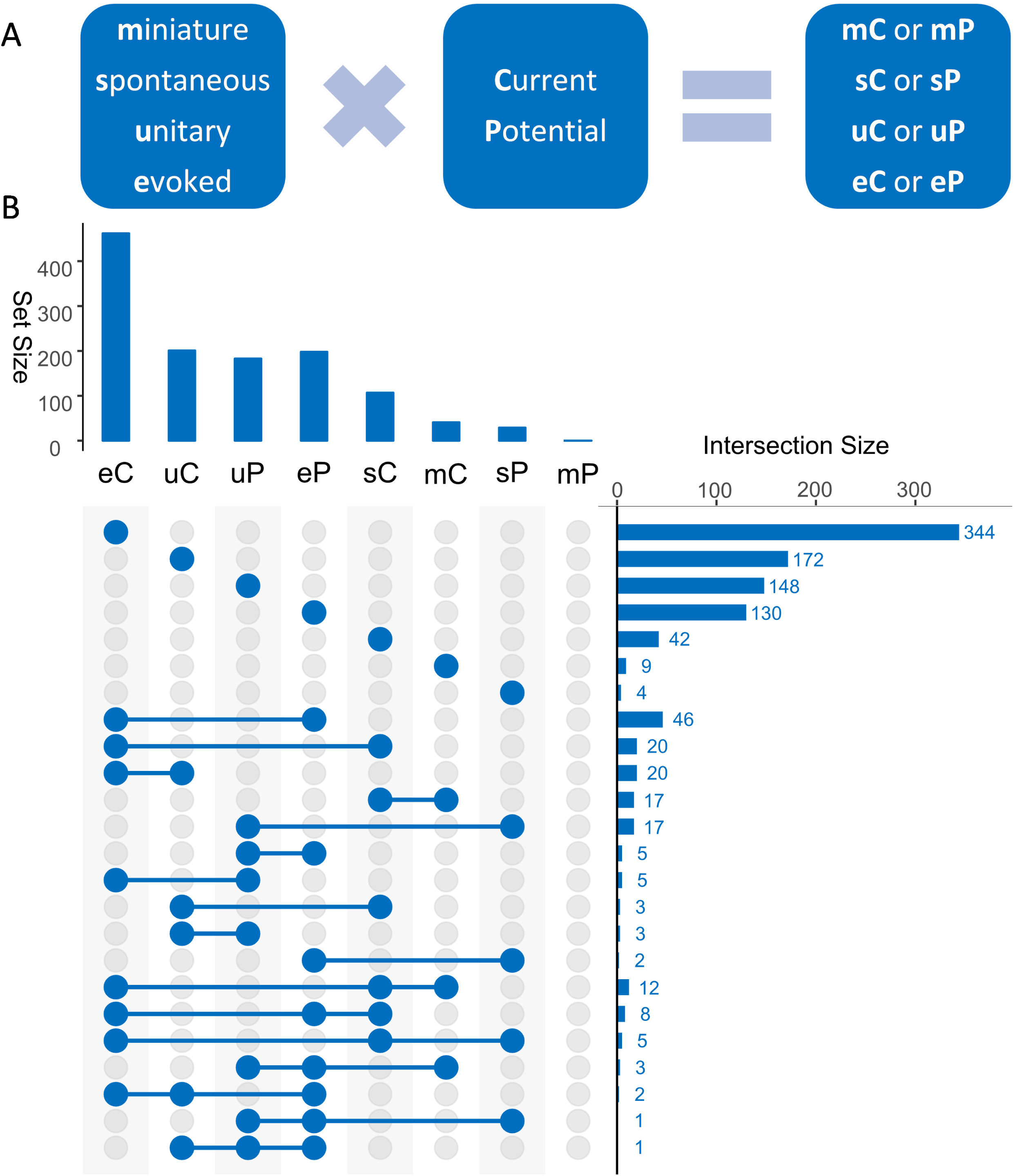
Data modalities. **(A)** Synaptic signals can be generated in eight different modalities depending on stimulation methods (e, u, s, and m) and response type (C or P). **(B)** The most prevalent modality among all extracted data (upper chart) is eC and the most prevalent combination of multiple modalities in the same experiment (right chart) is between eC and eP.

Besides the data modality distinction, the measurement methods and definitions to quantify the main properties of synaptic signals constitute another major source of data diversity compounding two distinct challenges. On the one hand, researchers characterize distinct aspects of amplitude, kinetics, and plasticity with complementary measures that are inherently incomparable. On the other, the terminology used to describe these measurements in scientific reports is itself ambiguous. As a result, even a relatively clear concept as “amplitude” may become difficult to relate between two studies of the same synapse type, because, when averaging signal traces, some researchers include all events, while others omit transmission failures. Some, but not all, reports refer to the latter case as “synaptic potency;” we adopt this nomenclature in the knowledge base to minimize confusion, but we always pay special attention to the correct identification of different measures by the reported definitions rather than the chosen name.

The situation is substantially more complex when extracting data pertaining to other synaptic features. Kinetics, for instance, can be characterized in terms of latency, rise, and decay time. Rise time can be reported as an exponential constant or as the temporal interval elapsed from 20% to 80% of the amplitude value among several other possibilities. Often rise and decay are combined as when reporting half-height signal width. With respect to plasticity, even if solely focusing on short-term dynamics and foregoing long-term changes, different experimental protocols may induce facilitation or depression and changes can refer to signal slope or amplitude, just to mention two of the many dimensions to consider. Since the terminology is here, too, inconsistent across studies, we have adopted the naming convention used by most studies, but employed prefixes and suffixes to differentiate conflicting names. Overall, besides synonymy, we have identified 68 actually distinct synaptic property measures (Fig. 5).

**Figure 5.**
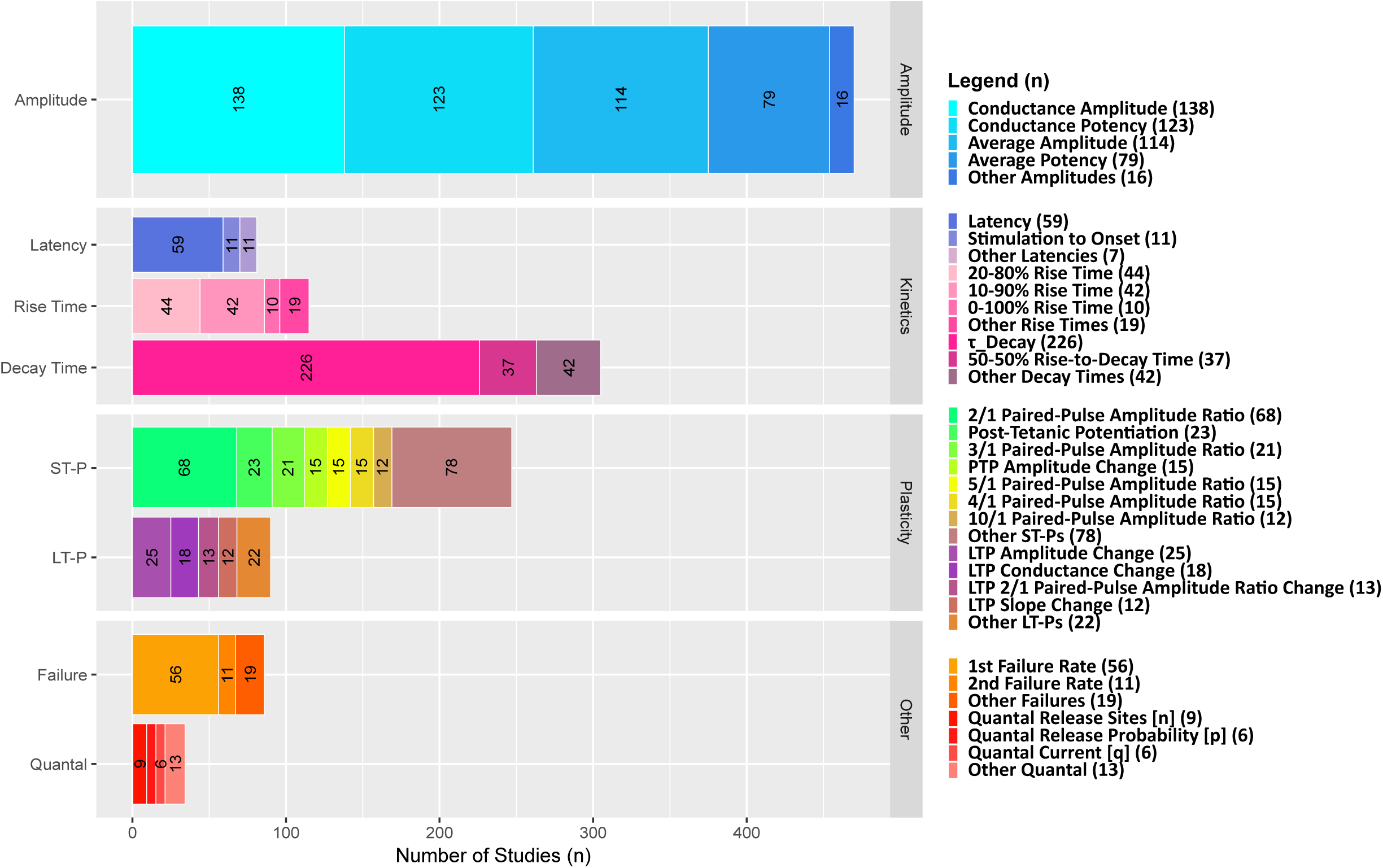
Measurement methods diversity. Synaptic conductance is the most prevalent measure for amplitude, single-exponential decay time constant (τ_Decay) for kinetics, paired-pulse ratio of 2^nd^ synaptic signal to the 1^st^ (2/1 Amplitude Ratio) for plasticity, and failure rate of the 1^st^ signal for other features.

Altogether, we have extracted 39,522 data entities: 8,486 (21%) constituting PoEs and 31,036 (79%) metadata (Table 2). Although paired recordings constituted approximately one quarter of mined experiments, they generated nearly 45% of the quantifiable synaptic evidence. Considering the diversity of stimulation protocols, recording modalities, and measurement types, the synaptic data are distributed across 619 columns in the master table of the knowledge base. The entire data collection is publicly released at Hippocapome.org/synaptome (Fig. 6). Users can search or sort synapse types based on the properties of pre- and post-synaptic neuron types. Each synapse type is linked to a list of experiment IDs. By following the experiment IDs on the “Evidence” tab, the user can access experiment summaries, annotated excerpts, and digitized traces. The same experiment IDs on the “Synaptic Data” tab provide access to the corresponding extracted values for amplitude, kinetics, plasticity, and other available properties.

**Table 2:** Pieces of evidence.

| Parameter | Amplitude | Kinetics |  |  | Plasticity |  | Other |  | Total |
| --- | --- | --- | --- | --- | --- | --- | --- | --- | --- |
|  |  | Latency | Rise time | Decay time | ST-P | LT-P | Failure rates | Quantal |  |
| PoE | 2,474<br>(29%) | 365<br>(4%) | 437<br>(5%) | 1,719<br>(20%) | 2,686<br>(32%) | 332<br>(4%) | 312<br>(4%) | 114<br>(1%) | 8,439<br>(100%) |

**Figure 6.**
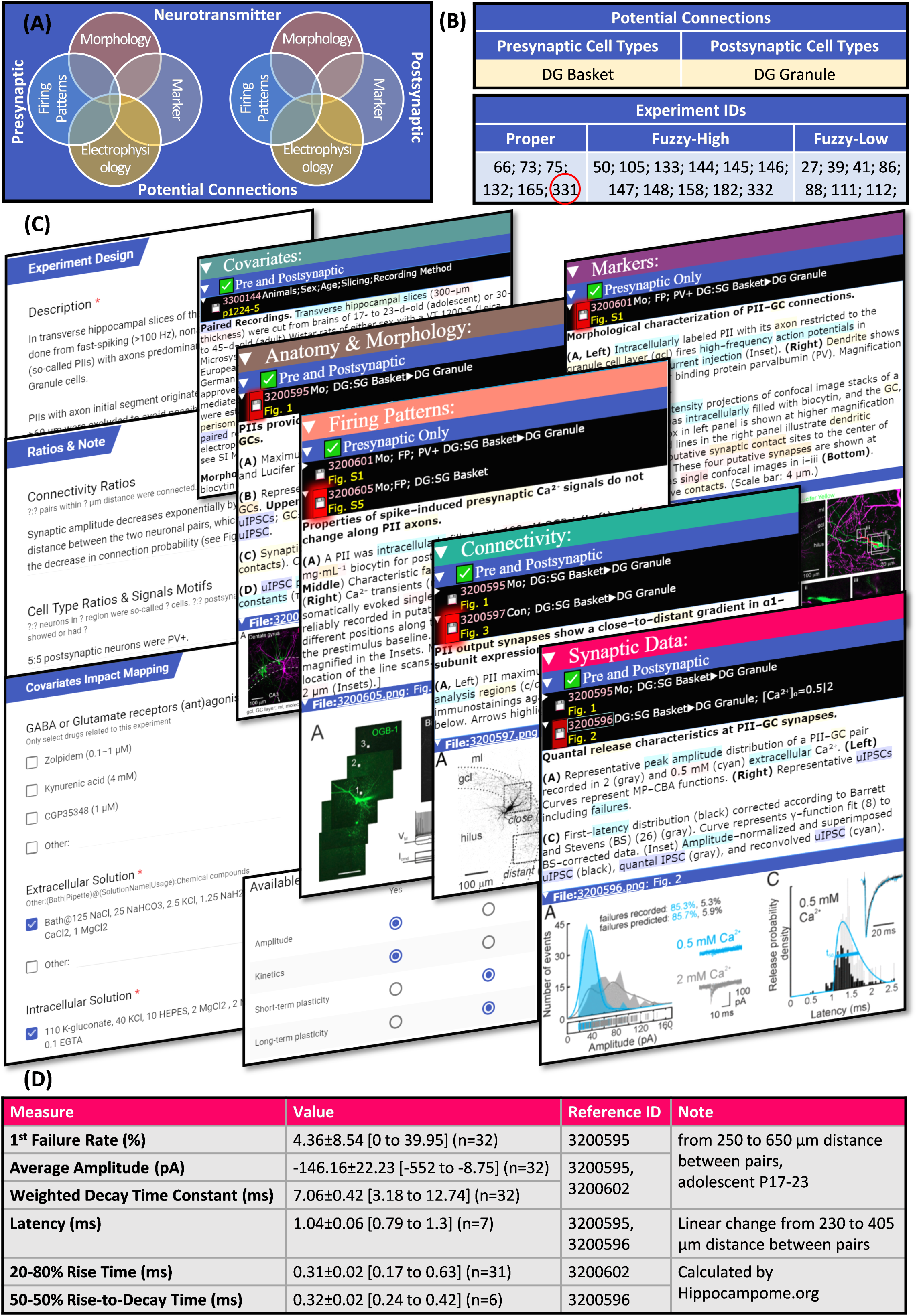
Data access. The described knowledge base of synaptic electrophysiology is freely available online. **(A)** Synapse types are searchable by the properties of the presynaptic and postsynaptic neuron types. **(B)** They are linked to experiment IDs categorized by mapping confidence. **(C)** The details and summaries of any experiment (for example, experiment with eID 331) can be reviewed while checking excerpts as evidence and **(D)** the extracted data is directly accessible. This example is from (Struber, Jonas, & Bartos, 2015).

### Data Integration and Usage Examples

The multiple sources and diverse settings of mined data make data normalization a necessity for meaningful comparisons. The junction potential *V*_j_, for instance, adds a systematic error that is usually not considered when reporting synaptic signal values. We have corrected all synaptic measurements based on the ionic composition of the interfacing solutions using our liquid-junction potential calculator, which is uniquely custom-designed to properly handle all recording methods and modes. Even after *V*_j_ correction, synaptic amplitude strongly depends on failure rate as well as on the difference between membrane and reversal potentials. To account for failure rate, we convert amplitude to potency by dividing the reported value by the success probability. Because electrophysiological studies are performed at widely different membrane potentials, data must be further normalized to conductance using Ohm’s law. This conversion, however, also requires a suitable *E*_rev_ value, which is not always measured or reported. The broad ranges of solution compositions used experimentally impose substantial differences in reversal potential. Thus, we calculate *E*_rev_ from the ionic composition of the bath and pipette solutions, recording method, and temperature.

With this normalization process in place, we used the subset of experimentally measured *E*_rev_ data to test the hypothesis that correcting for liquid junction potential would reduce the difference between experimental and theoretical reversal potentials. Remarkably, the *V*_j_ correction improved the agreement between calculated and measured *E*_rev_ for the vast majority of experiments (Fig. 7). Specifically, the average mismatch became statistically nil for both excitatory and inhibitory synapses in whole cell recordings as well as for GABA_A_ synapses recorded with sharp electrodes. For the limited sample of sharp-recorded AMPA synapses, the correction was neutral, possibly due to a larger margin of non-systematic error in this modality.

**Figure 7.**
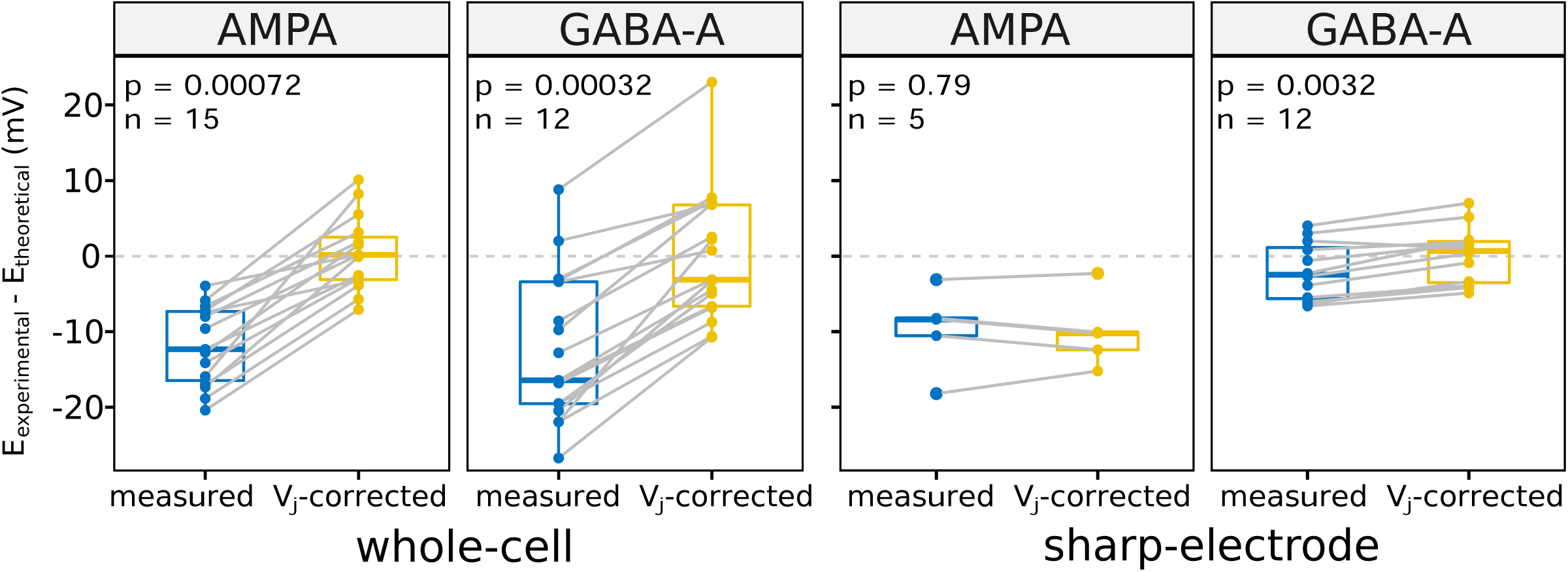
Correcting the liquid junction potential reduces the measured synaptic reversal potential error. After correcting the experimentally measured synaptic reversal potential (*E*) for liquid junction potential (*V*_j_), the difference between *E*_theoretical_ and *E*_experimental_ becomes close to zero on average. All data needed to calculate a pair (each grey line) come from one experiment with different solutions and temperature, which lead to different *V*_j_ and *E*.

Integrating reported synaptic data for thousands of synapse types provides a unique meta-analysis opportunity to confirm earlier observations and foster new discoveries. To this aim, we measured the correlation between specific properties of unitary GABAergic currents and different covariates (Table 3). In accordance with previous hippocampal studies, increasing the temperature reduces synaptic latency, decay kinetics, and paired-pulse ratio, making synapses faster, shorter, and more prone to show short-term depression than to short-term facilitation. Moreover, rising intracellular chloride concentration increases synaptic conductance and adding extracellular calcium or removing extracellular magnesium abates the synaptic failure rate. However, other previously suggested interactions did not reach statistical significance, such as the effects of holding potential or intracellular chloride concentration on decay time.

**Table 3:**
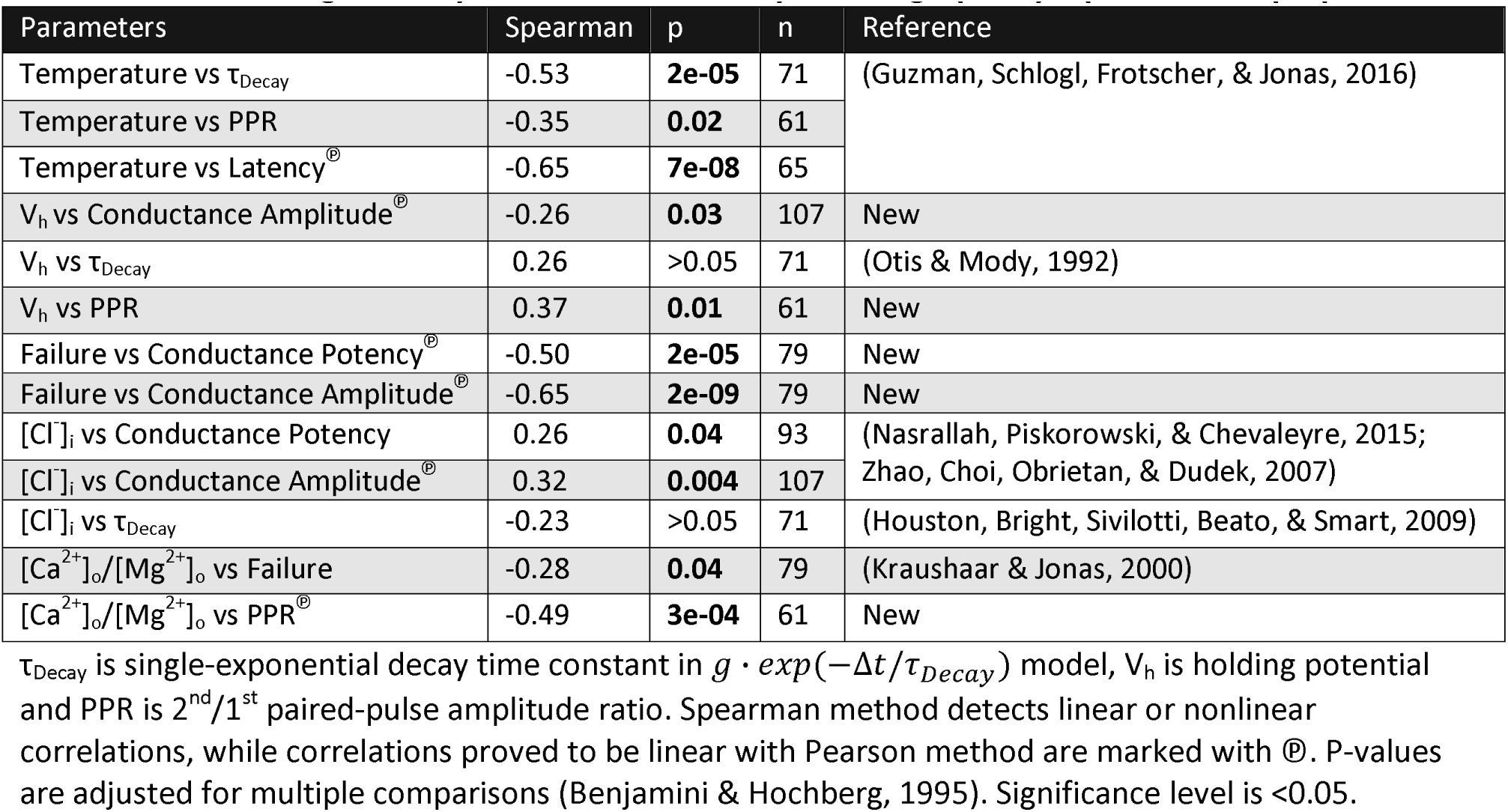
Covariates significantly correlate with unitary GABAergic postsynaptic currents properties.

We have also identified several relations that, to the best of our knowledge, had not been proposed before in the hippocampus literature. In particular, the paired-pulse ratio increases with postsynaptic membrane depolarization, but decreases with the extracellular concentration ratio between calcium and magnesium. Furthermore, synapses with higher failure rate tend to have smaller synaptic conductance potency. Last but not least, we checked whether faster GABAergic synapses also tend to be stronger and vice versa. The available data from unitary response indeed support a negative correlation between conductance potency and temporal decay (Fig. 8). Splitting the signals by decay time constant median accordingly results in a significant group difference in synaptic strength. Both the linear dependence and the effect size are more demarcated at physiological temperature than at room temperature.

**Figure 8.**
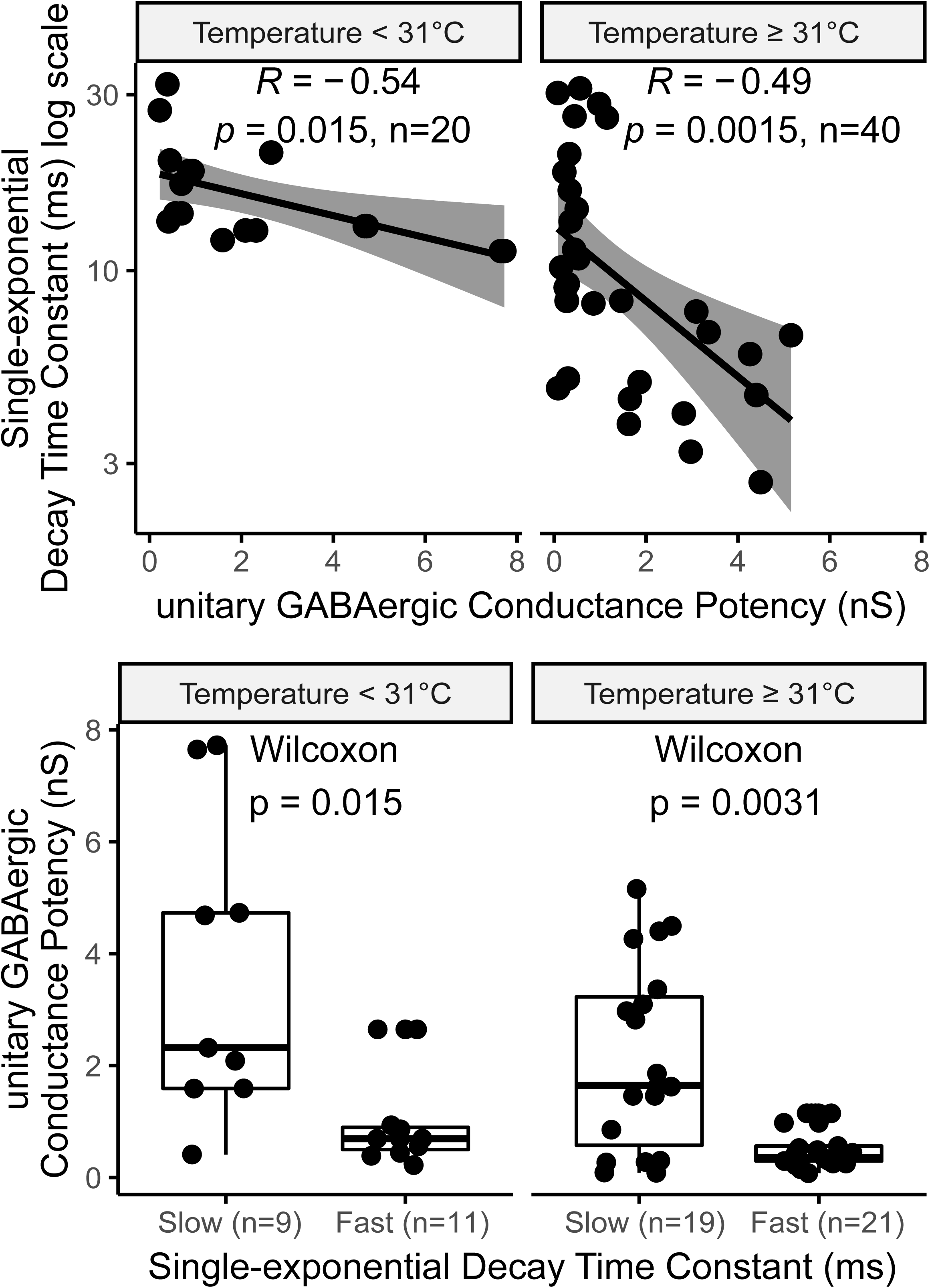
Faster GABAergic synapses are stronger. **(A)** GABAergic unitary synaptic potency significantly correlates with decay, both at temperatures ≥31°C or <31°C. **(B)** The conductance of slower synapses (decay time constant above median) is significantly smaller than that of faster ones. Each data point is the average or single result of a separate experiment.

## Discussion

Synapses play a crucial role in neural computation. Constraining large-scale biologically realistic simulations of brain circuits require detailed estimations of quantitative synaptic electrophysiology parameters (Bezaire, Raikov, Burk, Vyas, & Soltesz, 2016; Markram et al., 2015). Brain-wide catalogs of synaptic information may allow for data mining and systematic hypothesis testing. A recent high-throuput genetic imaging study in the mouse, for instance, ascribed a high degree of synaptic diversity to the hippocampal formation, possibly associated with its specific cognitive functions (F. Zhu et al., 2018). However, efforts in this direction have so far largely focused on molecular features (O’Rourke, Weiler, Micheva, & Smith, 2012; Paul et al., 2017; Zhang et al., 2007). The electrophysiological knowledge base of synapse types we have introduced and released with this report is the first of its kind for the rodent hippocampus or, to the best of our knowledge, any other neural system. Mining the data of more than 1,200 papers, we have extracted synaptic measures (amplitude and kinetics almost always, but also plasticity or other characteristics in most cases) or traces for 88% of synapse types, with substantially uniform coverage within and across the main sub-regions: dentate gyrus, CA3, CA1, and entorhinal cortex. Based on our comprehensive and methodical literature coverage, we surmise that the remaining synapse types have not yet been experimentally investigated and more research is therefore needed to ascertain their properties.

Continuing and extending the Hippocampome.org standard, we have associated all mined information to publication excerpts at the data-entity level. Therefore, anyone can immediately inspect the experimental context online and critically verify synapse type assignments, extracted data, and any corresponding assumptions or interpretations. We hope that such high level of transparency and provenance tracking will encourage constructive user feedback, allowing refinements and improvements in future releases. Importantly, this public resource also integrates experimental data produced with different methods, linking them together at the synapse-type level. Explicitly annotating experimental settings, recording modalities and covariates greatly facilitates the review of earlier works at single neuron-type detail, a feature no current search engine or automated text-mining approach provide. In addition, since we distinguish specific measurement types on the basis of their reported definitions (as opposed to adopted terminology), all data we group under one column of the accompanying database are guaranteed to be uniform, ensuring meaningful comparability across studies.

This opens the prospect of complete data normalization to enable quantitative data analysis by accounting for the effect of different covariates like temperature and other experimental metadata (Tebaykin et al., 2018). In theory, computational modeling should allow one to reduce the (at least) 68 distinct measures the hippocampal literature offers to quantify the eight possible recording and stimulation combinations into a handful of phenomenological parameters: one for amplitude, one for kinetics, and two or three for short-term plasticity (Tsodyks & Markram, 1997). As a start, we have calculated the reversal potential and corrected the systematic liquid junction error for all experiments. In the future, the correlations we have identified among specific synaptic measures and different covariates may also become useful for data normalization.

The majority of synaptic signals are recorded in slices. Extrapolating these in vitro manipulations to in vivo brain function is especially challenging because severed neuronal processes during sectioning alter connection probability and neuronal reconstruction integrity. For instance, one half of CA1 stratum oriens or stratum pyramidale neurons are partially axotomized in typical electrophysiological preparations, which affects morphological identification (Halasy, Buhl, Lorinczi, Tamas, & Somogyi, 1996; Kogo et al., 2004) and synaptic activity (Gulyas et al., 2010) differentially in longitudinal and transverse slices (Couey et al., 2013; Surmeli et al., 2015; Xiong, Metheny, Johnson, & Cohen, 2017). Homeostatic mechanisms also change synaptic strength after slicing in order to compensate for axonal loss (Dumas et al., 2018; Kirov, Sorra, & Harris, 1999), implying that the time elapsed since slice preparation (which is hardly ever reported in peer-reviewed publications) may be a key determinant of synaptic amplitude.

The choice of experimental method may even mask physiologically important synaptic properties. For instance, Cajal-Retzius cells’ unique cytosolic chemistry makes GABAergic input excitatory in these neuron types (Marchionni et al., 2010). Whole-cell recording not only changes intracellular ionic concentrations immediately after establishment, but also washes the signaling molecules responsible for synaptic plasticity (Bauer & LeDoux, 2004; Kato, Clifford, & Zorumski, 1993; Lamsa, Heeroma, & Kullmann, 2005; Maccaferri & McBain, 1996), an effect likely depending on cell size and synaptic distance from soma. Moreover, common use of gluconate in the pipette solution may change membrane electrophysiology, firing pattern, and even synaptic potentials in recorded neurons (Bullis, Jones, & Poolos, 2007; Fatima-Shad & Barry, 1993; Velumian, Zhang, Pennefather, & Carlen, 1997). Comprehensive data normalization and meaningful computer simulations will require a careful accounting of these effects.

In order to place this knowledge base content in appropriate circuit context, it is humbling to recognize the relatively low mapping resolution relayed through traditional literature reporting, with about 50% of experiments assigned to eight or more synapse types. Part of this degeneracy may be attributed to the existence of multiple morphological variants for certain neuron types; for example, while the canonical form of CA1 oriens/lacunosum-moleculare interneurons is characterized by an axonal tree ascending to lacunosum-moleculare, certain sub-types also branch off in radiatum or collateralize in oriens (Wheeler et al., 2015). Depending on the available experimental details, in many cases the ambiguity in neuron type identification is more consequential, as when equating all parvalbumin-expressing neurons to fast-spiking basket cells. To address these issues, our neuroinformatics pipeline translates all experiments in each study into machine-readable search queries, in order to find all corresponding potential connections automatically and in an unbiased way. Given the continuous evolution of modern technology (and of the very notion of neuron types in the neuroscience community), it is predictable that new cell types will be found and agreed upon, thus refining our understanding of hippocampal circuitry. Likewise, ongoing connectomics research will progressively validate and refute an increasing number of potential connections. Yet, the experimental information already reported in the scientific literature is not affected by these future changes and thus the resulting machine-readable queries will remain valid. Simply updating the neuron type definitions in Hippocampome.org will yield a revised mapping and, with additional neuron type properties identified, improved mapping resolution.

In the current experimental landscape, classic paired recordings produce the best mapping resolution and thus remains the most useful method to study synapses. In this method, both presynaptic and postsynaptic neuron types can be morphologically reconstructed while simultaneously measuring their molecular expression, membrane biophysics, and firing patterns. Furthermore, all eight different synaptic modalities are potentially recordable from the same neuron pair in one experimental setting. Although paired recordings can validate potential connections, it remains challenging to firmly refute potential connections electrophysiologically. Even with a large enough sample size to avoid sampling bias, most stimulation protocols may be unsuitable to rule out synaptic connectivity. For instance, the low initial release probability of certain hippocampal synapses requires 10 to 20 stimuli to trigger a detectable signal (Szabadics & Soltesz, 2009). An ideal experiment needs an adequate number of presynaptic stimuli at different frequencies. Optimizing the stimulation paradigm may be useful to conduct the maximum number of informative synaptic experiments in a short time neuron remain viable.

Increasing the number of electrodes, the connectivity of examinable neuronal pairs grows quadratically (Perin & Markram, 2013). For instance, octuple whole-cell recording enables the parallel investigation of up to 56 (8×7) neuron pairs, while examining evoked and background synaptic activity, morphology, electrophysiology, and biochemistry of eight neurons (Jiang et al., 2015). Automated robotic patch clamp can further augment the data yield (Bruggemann, Stoelzle, George, Behrends, & Fertig, 2006; Kodandaramaiah, Franzesi, Chow, Boyden, & Forest, 2012; Lepple-Wienhues, Ferlinz, Seeger, & Schafer, 2003; Vasilyev, Merrill, Iwanow, Dunlop, & Bowlby, 2006). Combining optogenetically-enabled photostimulation (Kim, Adhikari, & Deisseroth, 2017) or single-neuron genetic profiling by patch-seq (Cadwell et al., 2016) may eventually enable the compilation of a whole-brain, high-resolution functional and multimodal map of brain synapses.

Although effective for statistical analysis and widespread in reporting practice, data pooling lowers the mapping resolution of experiments (as illustrated in Fig. 2). To alleviate this problem and increase the impact of studies, we recommend the release of experimental data, at least in the form of supplemental tables, thus allowing meta-analysis and data re-usage. Each synaptic measure should be linked to the morphology, biomarkers, electrophysiology, and firing patterns of the corresponding neuron pair. Researchers can also use our framework to organize their data, which optimizes the mapping, usage, and visibility of their data and prevents the information loss inherent in within-study data pooling.

In contrast, collating datasets across laboratories, experimental techniques, geographical locations, publication years, and animal subjects may be a powerful approach for discovering general trends, anomalous results or interesting covariates. For example, our meta-analysis revealed several new correlations that would have been impossible to detect from a single source and a uniform set of metadata. These findings are valuable and robust because of both larger sample size and data source diversity, allowing the discovery of truly invariant synaptic properties. Moreover, having collected and organized all available electrophysiology data concerning hippocampal synapses will facilitate the construction of better computational models of this important neural system while helping researchers find gaps in scientific knowledge and compare new data with existing ones. We also hope this public resource will encourage multidisciplinary approaches to complex multimodal neurobiological data interpretation.

## Acknowledgments

The authors are grateful to Diek Wheeler, Christopher Rees, David Hamilton, Carolina Tecuatl, Siva Venkadesh, Amar Gawade, Grey Madison, Zainab Aldarraji, Patricia Maraver, and the Hippocampome.org summer interns for their contribution and feedbacks. This research was supported in part by George Mason University’s Provost Office through a Dissertation Completion award as well as by National Institutes of Health through grant U01MH114829 (BICCN) and R01NS39600 (BISTI).

